# Brain signatures indexing variation in internal processing during perceptual decision-making

**DOI:** 10.1101/2023.01.10.523502

**Authors:** Johan Nakuci, Jason Samaha, Dobromir Rahnev

## Abstract

Brain activity is highly variable even while performing the same cognitive task with consequences for performance. Discovering, characterizing, and linking variability in brain activity to internal processes has primarily relied on experimentally inducing changes (e.g., via attention manipulation) to identify neuronal and behavioral consequences or studying spontaneous changes in ongoing brain dynamics. However, changes in internal processing could arise from many factors, such as variation in strategy or arousal, that are independent of experimental conditions. Here we utilize a data-driven clustering method based on modularity-maximation to identify consistent spatial-temporal EEG activity patterns across individual trials and relate this activity to behavioral performance. Subjects (N = 25) performed a motion direction discrimination task with six interleaved levels of motion coherence. Modularity-maximization based clustering identified two discrete spatial-temporal clusters, or subtypes, of trials with different patterns of brain activity. Surprisingly, even though Subtype 1 occurred more frequently with lower motion coherence, it was nonetheless associated with faster response times. Computational modeling suggests that Subtype 1 was characterized by a lower threshold for reaching a decision. These results highlight trial-to-trial variability in decision processes usually masked to experimenters and provide a method for identifying endogenous brain state variability relevant to cognition and behavior.

**Highlights:**

⍰ Brain activity is highly variable.
⍰ We find multiple and distinct stimulus-driven patterns in EEG.
⍰ With changes in decision-making and drift-diffusion parameters.
⍰ These results suggest a new way to identify brain states relevant to behavior.

**Graphical Abstract:** 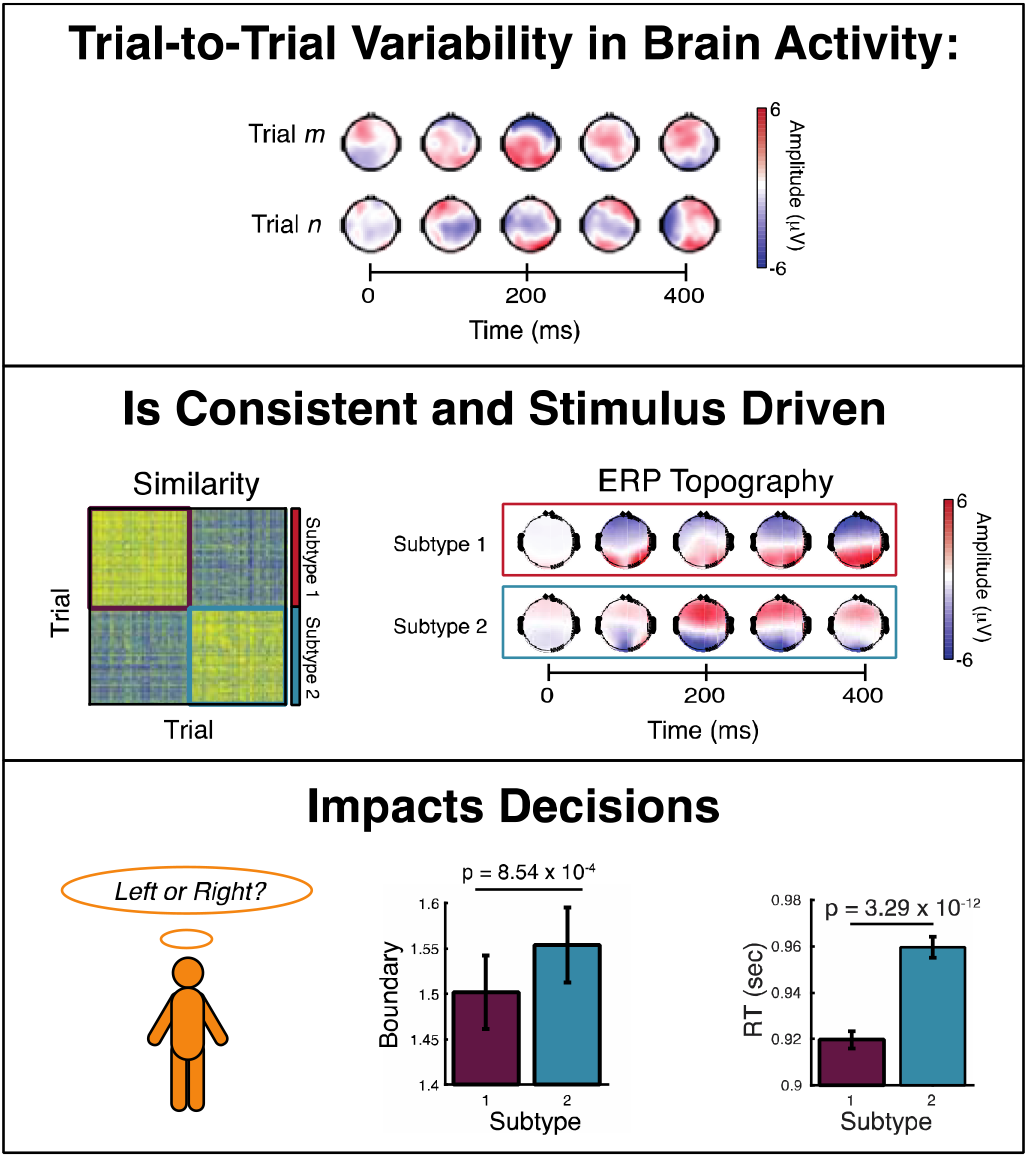

## Introduction

Variability in performance is present in many day-to-day activities and cognitive tasks^1–7^. Alongside variability in performance, brain activity during a task is highly variable^1,4^ and may impact the efficiency and accuracy of task performance^8–11^. Trial-to-trial variability in brain activity is present at all levels of cortical organization from individual neurons^12^ to large-scale brain networks^13^. This variation in brain activity impacts cognition and behavior in social situations^14^, economic decisions^15^, and in low-level perception^16^. Despite the widespread variability in brain activity during a task, standard analyses aim to identify event-related changes in brain activity that are consistent across all trials^17^.

When measuring brain activity with Electroencephalography (EEG), stimulus-driven activity can be identified with event-related potentials (ERPs), which provide the average time-locked response to an experimental event^18^. However, it is also possible that subsets of trials exhibit brain activity that is not well represented by the average^19,20^. Indeed, brain activity measured with EEG is both spatially and temporally variable during a task, with at least some of this variability likely stemming from meaningful variation in internal processing rather than simply noise^8–11^. Additionally, averaging across all trials might eliminate the ability to observe distinct forms of stimulus-driven activity (i.e., the ERP) that are engaged differentially on individual trials or contexts but that are, nonetheless, relevant for behavioral performance.

Here, we link trial-to-trial variation in brain activity measured with EEG and decision-making processes during a perceptual decision-making task using a data-driven classification method we developed previously^19^. Briefly, modularity-maximization identified unique stimulus-driven brain activity variation among repeated trials of a motion discrimination task^21^. To anticipate our results, modularity-maximation identified two distinct subtypes of stimulus-driven brain activity. Further, we establish the behavioral profile associated with each subtype and link this variation to underlying latent cognitive processes. Overall, our results indicate that multiple brain activity patterns can co-exist in the context of the same task.

## Results

Here, we identify multiple consistent patterns of stimulus-driven activity among individual trials and link this variation to behavior and underlying latent cognitive processes. Subjects performed a motion discrimination task where they judged the global direction of a set of moving dots (left/right) with six levels of coherence (**Figure 1A**). Even in a simple and repetitive task such as this, trial-to-trial spatial and temporal variation in brain activity measured with EEG is evident (**Figure 1B**).

**Figure 1.**
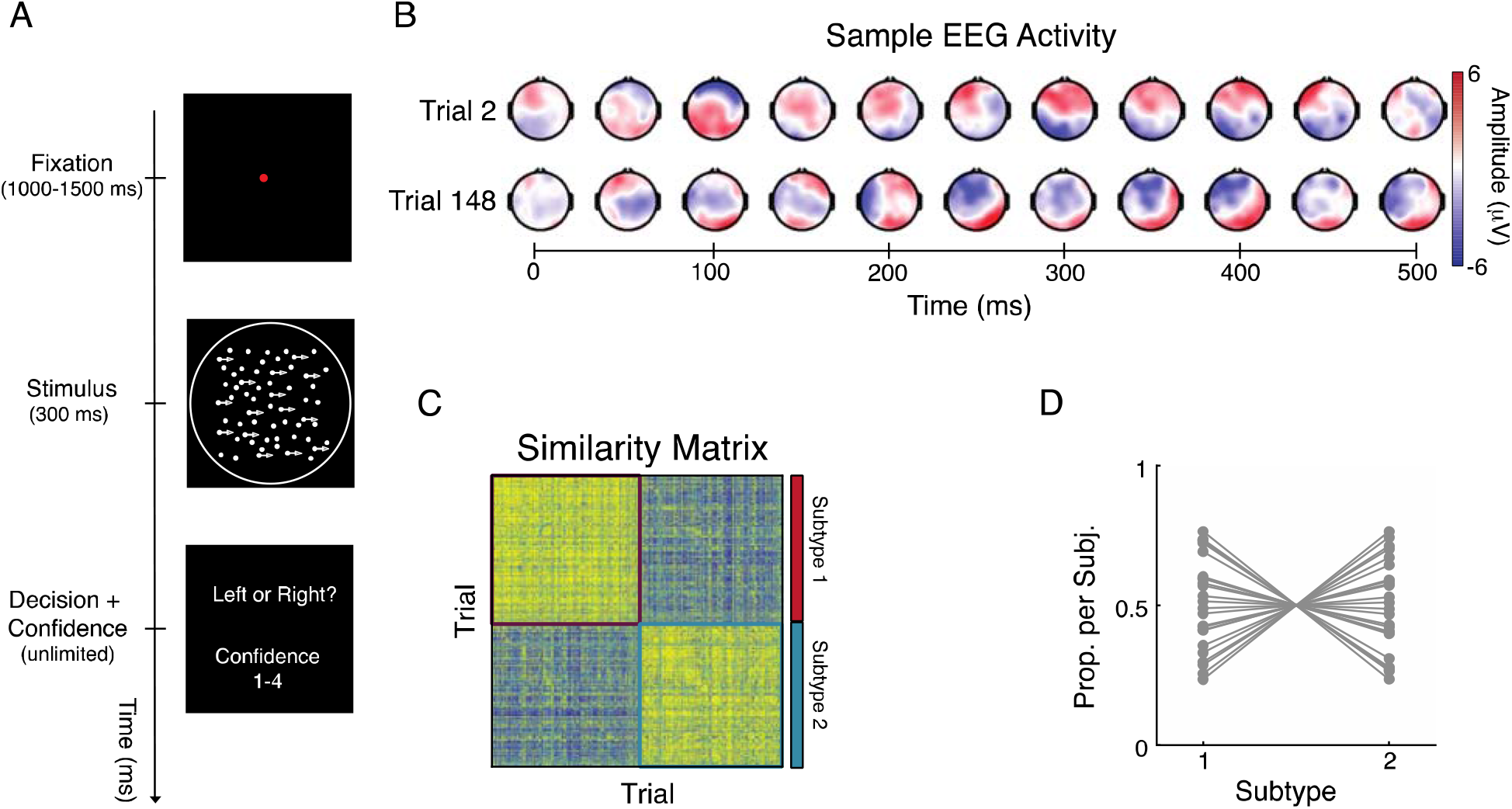
Subtypes of individual trials in a motion perception task. A) Subjects viewed a dot motion stimulus for 300 ms with net motion direction either to the left or the right at varying levels of motion coherence (arrowed dots). Using a single button press, subjects provided a choice and confidence (1-4) judgment. B) EEG activity from two trials from stimulus onset (0 ms) to 500 ms after onset from the same subject. The brain activity between the trials exhibits stark differences. C) Modularity-maximization based clustering identified two subtypes of trials, Subtype 1 and Subtype 2. The colored squares correspond to the trials composing each subtype. Pearson correlation was used to calculate the spatial-temporal similarity of the EEG activity among individual trials from 0 to 500 ms post-stimulus. D) The proportion of trials in each subject classified as either subtype 1 or 2.

To identify consistent patterns in brain activity across trials, we pooled all trials across subjects together to calculate the spatial and temporal similarity using Pearson correlation from stimulus onset (0 ms) to 500 ms after onset. The modularity-maximization classification procedure identified two subgroups of trials, Subtype 1, which accounted for 50.87% of trials (N_trials_ = 10674), and Subtype 2 accounted for 49.01% of trials (N_trials_ = 10284; **Figure 1C**), across all subjects (**Figure 1D**).

To understand the nature of the two subtypes, we plotted their average event-related potentials (ERPs) to test for differences in stimulus-driven activity^18^. Qualitatively, the ERPs for each subtype exhibited an opposite pattern of anterior vs. posterior event-related potentials (**Figure 2A**). These qualitative topographical differences were present even when comparing ERPs for each motion coherence level (**Figure S1A, B).** To confirm these impressions, we compared the topographical similarity of ERPs estimated for each subtype using a topographic analysis of variance (TANOVA)^22^. We found strong and consistent differences between subtypes in the post-stimulus period (p < 0.001, FDR corrected; **Figure 2A**). Further, focusing on the centro-parietal sensor, which has been linked with decision-making processes^23^ and evidence accumulation^24–26^, significant differences were present in amplitude between the subtypes (independent samples t-tests: p < 0.001, FDR corrected; **Figure 2B**) and for each motion coherence level (independent samples t-tests: p < 0.001, FDR corrected; **Figure S1C**). Subtype 1 contained significant positive amplitude in the parietal area compared to Subtype 2 from stimulus onset (0 ms) to 1000 ms, extending beyond the 500 ms window used in the clustering.

**Figure 2.**
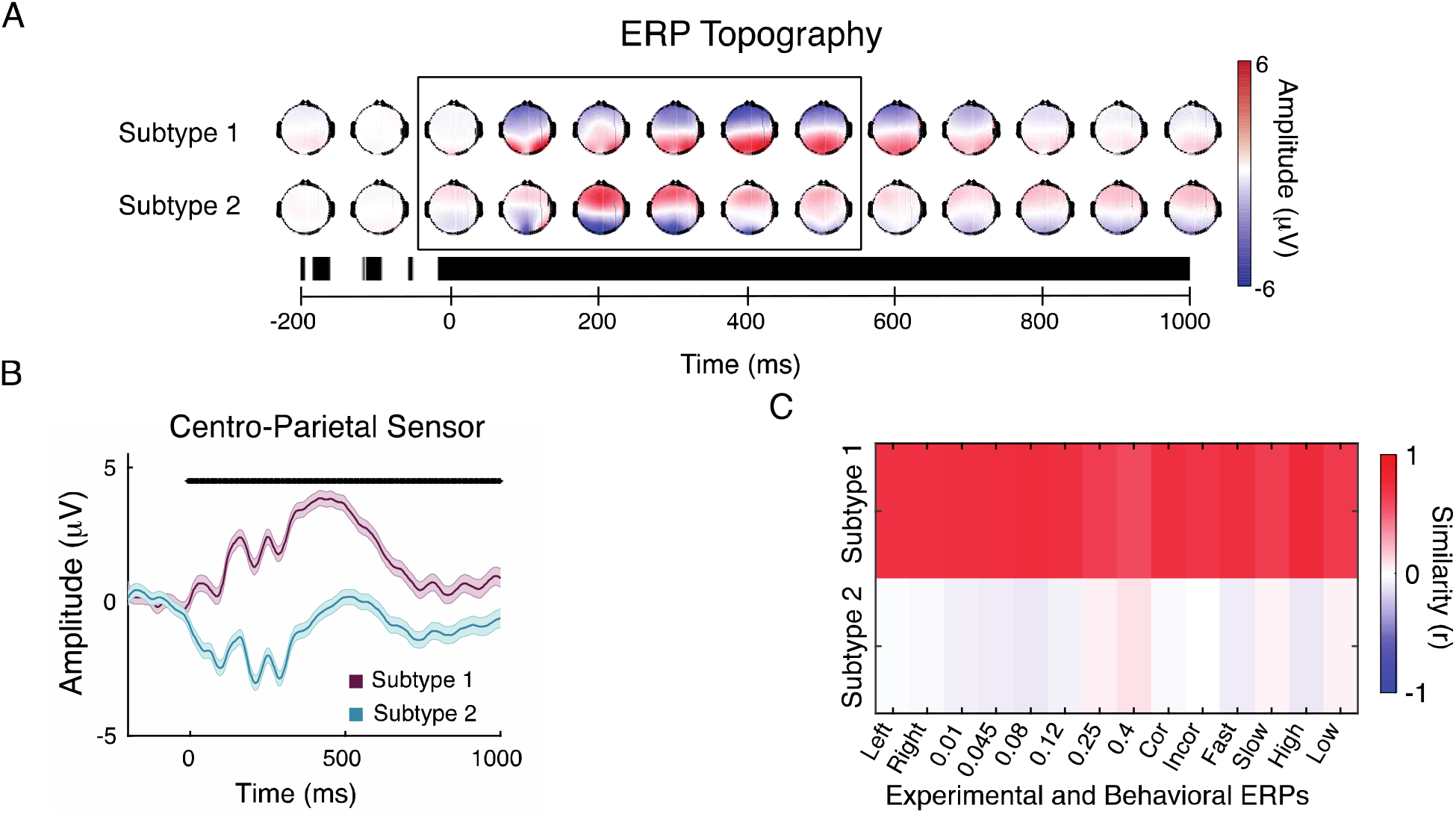
Differential patterns of stimulus-driven brain activity between subtypes. A) ERP topographies of Subtype 1 and Subtype 2 from 200 ms before stimulus onset to 1000 ms after stimulus offset (*top*). Note that the clustering algorithm was applied to the data from stimulus onset (0 ms) to 500 ms, black box. Topographical ANOVA identified periods of topographic differences between subtypes. Black regions indicated periods where the differences were significant (*bottom*). B) ERP activity from the centro-parietal sensor per subtype. Each waveform shows the mean (thick line) and standardized measurement error of the mean (shaded area)^30^. Statistical testing was conducted using independent samples t-tests, and FDR corrected for multiple comparisons. Statistically significant differences in amplitude are marked at the top of the panel. C) The topographical similarity between subtype-derived ERPs to ERPs derived from experimental – motion direction (Left/Right), motion coherence (0.01, 0.045, 0.08, 0.12, 0.25 0.4) – and behavior factors – Correct/Incorrect response, Fast/Slow response time, High/Low confidence. Pearson correlation was used to calculate the spatial-temporal similarity of the EEG activity from 0ms to 1000 ms after the stimulus. The ERP from one of the subtypes, Subtype 1, exhibits strong similarity (r > 0.60) to ERPs derived from experimental and behavioral factors highlighting the utility of Modularity-Maximization based clustering to identify variation in internal processing relevant to cognition.

One possibility is that these subtypes reflect ERPs associated with different experimental or behavioral factors, such as leftward/rightward moving trials or fast/slow responses. To better assess the nature of these subtypes, we compared the topographical similarity between subtype-derived ERPs to ERPs derived by averaging trials associated with experimental (motion direction and coherence levels) and behavioral (accuracy, response times, and confidence) factors. The topographical similarity was estimated between ERPs from stimulus onset (0 ms) to 1000 ms after. Interestingly, a strong similarity was found in Subtype 1 (r > 0.60) but not in Subtype 2 (r < 0.10) to ERPs derived from experimental and behavioral factors, indicating that in 49.01% of trials from our study, the variation in the stimulus-locked ERP was induced by other factors (**Figure 2C**).

Additionally, we tested if subtypes were associated with differences in pre-stimulus brain activity. Pre-stimulus brain activity has been found to impact post-stimulus brain activity and behavioral performance^27–30^. To investigate differences in the pre-stimulus brain activity, we tested for differences in power from -200 ms to 0 ms in the delta, theta, alpha, and beta frequency bands (**Figure 3**). We found the strongest differences between the subtypes in alpha (paired-samples t-test, t(24) = 5.95, p = 3.8×10^-6^; **Figure 3C**) and beta bands (paired-samples t-test, t(24) = 4.79, p = 3.5×10^-5^; **Figure 3D**). These findings add to the growing body of literature showing that pre-stimulus activity affects post-stimulus processing^31–33^.

**Figure 3.**
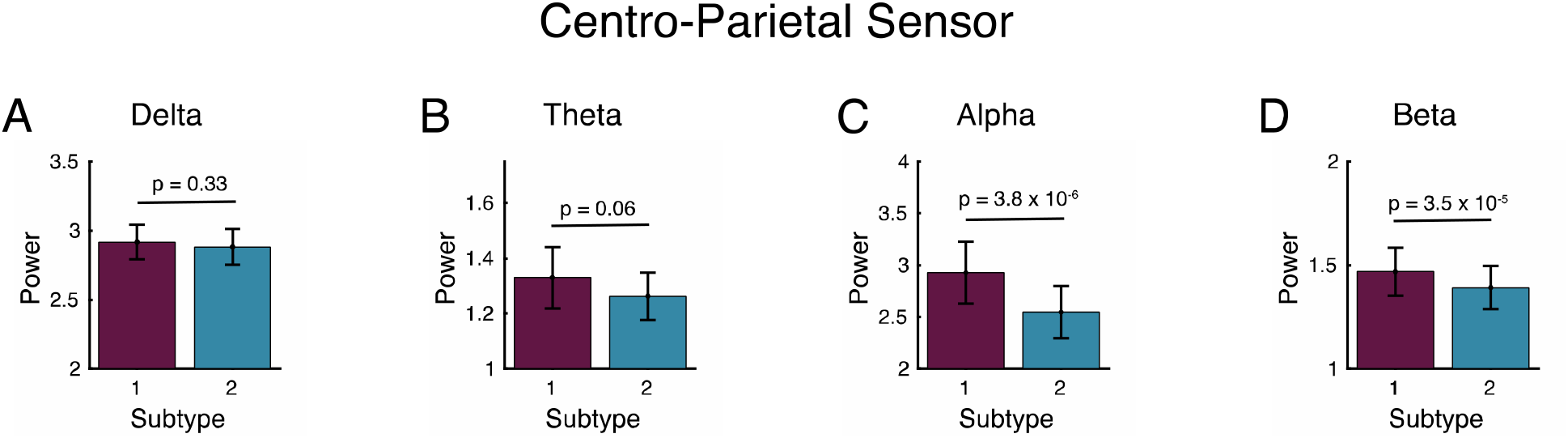
Differences in pre-stimulus brain activity between subtypes. Differences in average power from -200 ms to 0 ms for the A) delta (1-3 Hz), B) theta (4-7 Hz), C) alpha (8-12 Hz), and D) beta (13-30 Hz) bands. The average power was calculated for the centro-parietal sensor after first filtering the EEG signal and followed by Hilbert transformation. Error bars show the mean ± sem.

Importantly, we examined whether the subtypes reflect differences in the composition of trials based on experimental factors. We found that the distribution of trials with leftward and rightward motion was the same between subtypes (Wilcoxson rank sum test: Z = 0.13, p = 0.89; **Figure 4A**). However, Subtype 1 contained a higher proportion of trials with lower motion coherence (Wilcoxson rank sum test: Z = -4.06, p = 4.72×10^-5^; **Figure 4B**), this difference accounted for less than 3% of trials per condition (**Figure 4C)**. Thus, experimental factors were not the main driver of the spatial-temporal variation in brain activity among trials.

**Figure 4.**
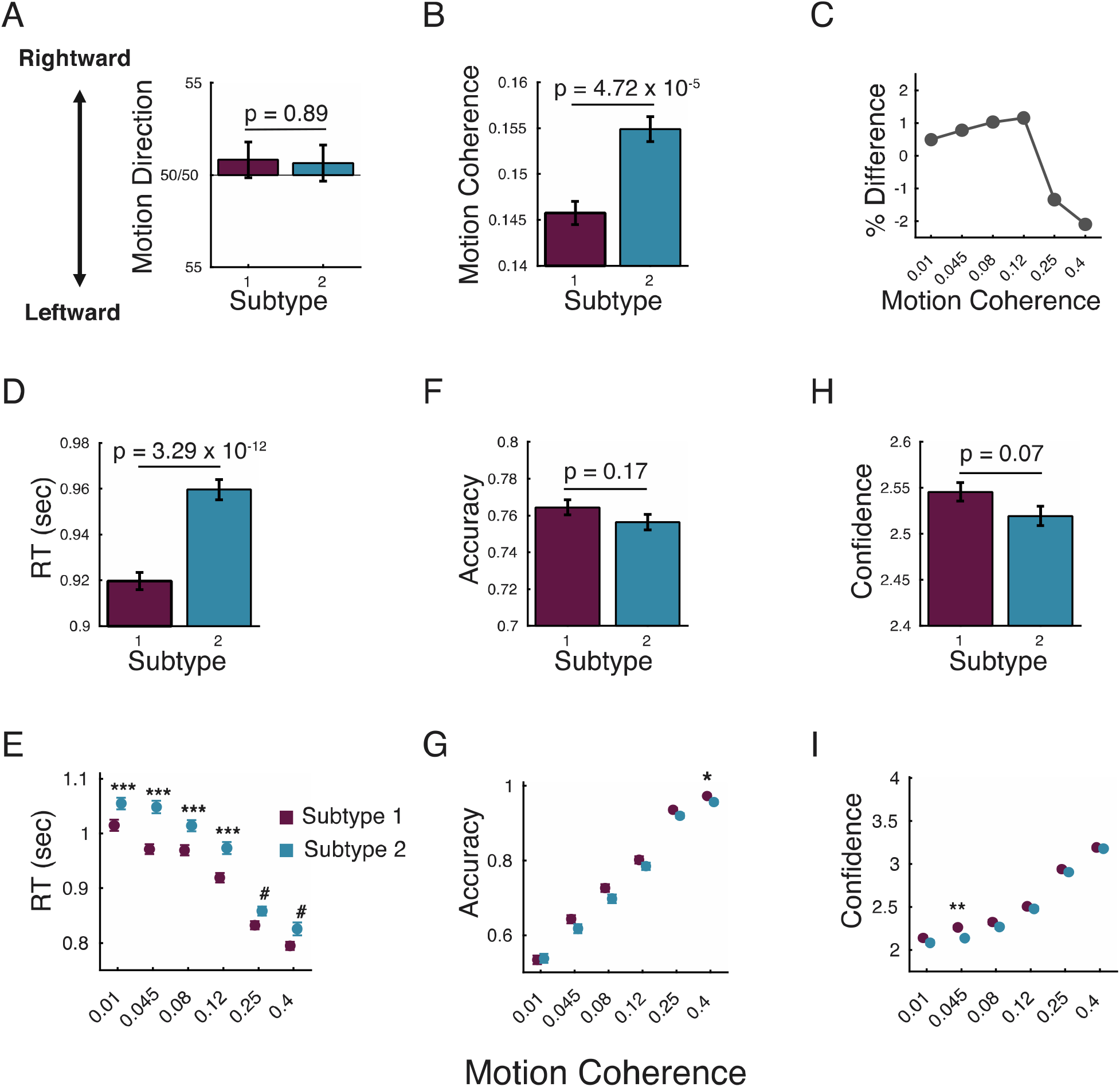
Experimental and behavioral differences between subtypes. Corresponding differences in (A) Motion Direction and (B) Motion Coherence level between subtypes. C) Differences in the percent of trials between subtypes per motion coherence level. Differences in (D-E) response times, (F-G) accuracy, and (H-I) confidence between subtypes. Error bars show the mean ± sem. *** p < 0.001, ** p < 0.01, * p <0.05 FDR corrected; # p < 0.05 uncorrected.

Further, we investigated the average number of consecutive trials (dwell time) for each subtype in order to test if the subtypes reflect slow processes spanning multiple trials. We compared the dwell times between Subtype 1 and Subtype 2 to a randomized re-ordering of subtype labels. We found no significant differences in dwell time between each of the comparisons (paired-samples t-test, p > 0.05; **Figure S2A**), suggesting that the two subtypes occur in a largely random fashion with no clear repeating patterns. Additionally, we examined if there were differences in the proportion of trials between the first and second half of the experiment and found that the proportion of Subtype 1 trials increased from the first to the second half of the experiment (paired-samples t-test, p < 0.01; **Figure S2B**), but the differences between the two halves were relatively small, 3.1% and 3.2% trial, respectively. These results suggest that the subtypes do not reflect associated slow processes spanning multiple trials or learning associated changes.

Critically, we investigated whether the subtypes reflect differences in behavioral performance. We utilized a mixed-effect model to assess the effects of the subtype, motion coherence and motion direction on reaction times, accuracy, and confidence. As expected, we found that increased motion coherence led to lower reaction time (t(20956) = -4.34, p = 1.40×10^-5^), higher accuracy (t(20956) = 6.54, p = 6.43×10^-11^), and higher confidence (t(20956) = 9.00, p = 2.44×10^-19^). Critically, the subtypes significantly differed in reaction time (t(20956) = 3.53, p = 4.5×10^-5^), but not accuracy (t(20956) = 1.02, p = 0.31) or confidence (t(20956) = -0.66, p = 0.50). The differences in reaction times between subtypes can be observed when comparing within each motion coherence level, Subtype 1 trials consistently exhibited faster response times (independent samples t-test: t(20956) = -6.97, p = 3.29×10^-12^; **Figure 4D, E)**. On the other hand, no significant differences were present between the two subtypes in accuracy (independent samples t-test: t(20956) = 1.35, p = 0.17; **Figure 4F, G)**, and only marginally higher confidence in Subtype 1 (independent samples t-test: t(20956) = 1.79, p = 0.07; **Figure 4H, I**).

Having identified two trial subtypes with underlying differences in stimulus-driven brain activity and decision-making processes, we sought to determine the latent cognitive processes that would give rise to the behavioral differences by computationally modeling the response times and accuracy using the drift-diffusion model^34^. We fit the drift-diffusion model to the behavioral data from each subtype separately. We let the drift rate vary with motion coherence level, but the decision boundary and non-decision time were fixed across the different coherence levels. The drift-diffusion model was able to reflect behavioral data quite well. The predicted reaction times for Subtype 1 were consistently faster (independent samples t-test; p < 0.05; **Figure 5A**) and with accuracy exhibiting no differences (independent samples t-test; p > 0.05; **Figure 5B**).

**Figure 5.**
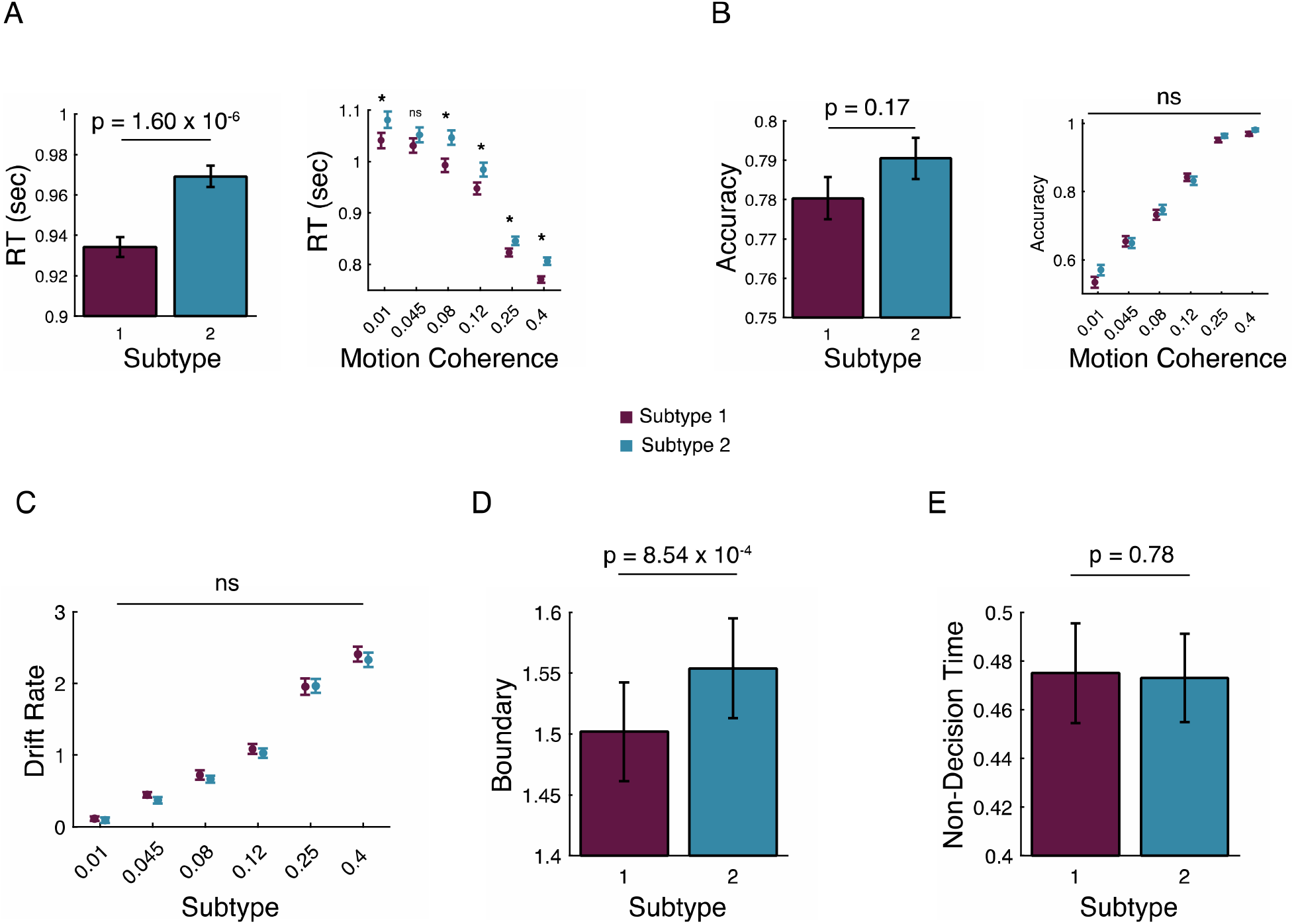
Drift-diffusion model. Drift-diffusion modeling results reflected behavioral performance with (A) reaction times being faster in Subtype 1 and (B) no significant differences in accuracy. *Left* panels in A and B represent the average reaction times and accuracy. *Right* panels show the performance for each motion coherence level. Drift-diffusion parameters showed that (C) the drift rate was the same between subtypes, (D) the response boundary was higher in Subtype 2, and (E) the non-decision time exhibited no differences between subtypes. Statistical testing was conducted using independent samples t-tests, and FDR corrected for multiple comparisons. * p < 0.05; ns = not significant

Critically, examining the latent factors, we found the drift rate was the same between subtypes (paired-samples t-test: p > 0.05; **Figure 5C)**, but Subtype 2 trials had a significantly higher response boundary (paired-samples t-test: t(24) = -3.81, p = 0.001; **Figure 5D)**. Further, no differences were present in the non-decision time (paired-samples t-test: t(24) = 0.28, p = 0.81; **Figure 5E).**

Lastly, we conducted two additional analyses to ensure our results were generalizable and robust. First, we performed a 5-fold cross-validation analysis using Support Vector Machine (SVM). An SVM classifier correctly predicted subtype labels with greater than 98% accuracy (**Figure S3A**). Further, the classification weights were consistent across EEG channels and time, suggesting that the EEG activity used to separate the trials was spatially and temporally distributed (**Figure S3B-D**). Second, we replicated the analysis with a longer time clustering window (1000 ms) to verify that the results were not dependent on the time range used in the clustering analysis. The classification similarity between the 500 ms and 1000 ms time windows was strong (>84%; **Figure S4A-C)** which is reflected in the ERP and behavioral analysis (**Figure S4D-O)**.

## Discussion

Behavior and brain activity are highly variable. Behavioral variability is ubiquitous in social situations^14^, economic decisions^15^, and even low-level perception^16^. Variability in brain activity is present from the levels of individual neurons^12^ to large-scale brain networks^13^. However, identifying the underlying neuronal mechanism associated with behavioral variability has been challenging^35^. Nonetheless, variation in internal processing that impacts behavior still needs to be discovered and characterized^36^.

To address this issue, we utilized a data-driven clustering method based on modularity-maximation. The key component in the analysis is that individual trials are clustered based on their spatial-temporal similarity in the post-stimulus period to identify consistent patterns of brain activity. Here spatial-temporal similarity is estimated across sensors and time. We found that individual trials could be separated into clusters, which we called Subtype 1 and Subtype 2.

Critically, for each subtype we can determine the underlying stimulus-driven brain activity by averaging the trials in each subtype to estimate the ERP. The two subtypes contained ERPs with different spatial-temporal activity patterns and differed in reaction times. Further, we found stronger posterior alpha and beta power in the pre-stimulus period from -200 ms to stimulus onset in Subtype 1. Computational modeling indicated that differences in reaction time arose from alterations in the threshold for reaching a judgment. These results demonstrate that brain activity measured with EEG can be used to distinguish subtypes of trials differing in their underlying internal processes.

Traditionally, identifying variation in stimulus-driven internal processing has relied on examining predefined brain features such as EEG low-frequency power^37–39^ or the slope of the 1/f spectrum^40,41^. However, these approaches could limit the identification of brain activity patterns relevant to cognition because it limits the sources of neuronal activity that could be contributing to behavioral variability. Our data-driven analytical framework overcomes these limitations and was able to identify brain activity important for cognition and behavioral performance.

These two different subtypes could indicate the existence of different cognitive modes. Recent studies have suggested that humans^42^ and other animals^43^ switch between different modes of processing during perceptual decision-making tasks. These modes could arise from changes in a single information processing sequence induced by alteration in the balance between top-down^44^ and bottom-up signaling^45^. Alternatively, the different stimulus-driven activity could indicate the existence of two independent information processing sequences.

Lastly, it is important to highlight the limitations of our methodology and analysis. One limitation pertains to the specific set of decisions made regarding the clustering, such as the time window, trial-to-trial similarity estimation, and the clustering method. For example, phase-based measures of similarity instead of Pearson correlation might result in different clusters. Future research should investigate how different ways of clustering affect the results to determine the consistency of the clusters identified here. A second limitation pertains to the interpretation one can draw from our results. The analysis yielded two clusters, which can naturally be interpreted as indicating the existence of two discrete states. However, it is possible that the states are continuous instead of discrete, and only appear discrete due to the clustering in discrete bins. Future research should dissociate between these two possibilities.

In conclusion, we find two forms of stimulus-driven brain activity present in all subjects but exhibiting stark differences in topographical organization and affecting behavioral performance. These results have strong implications for the common practice of identifying stimulus-driven brain activity by averaging across trials such that the brain may contain multiple mechanisms for reaching a decision. The analytical approach and findings presented here open a new avenue for understanding the brain-behavior relationship.

## Methods

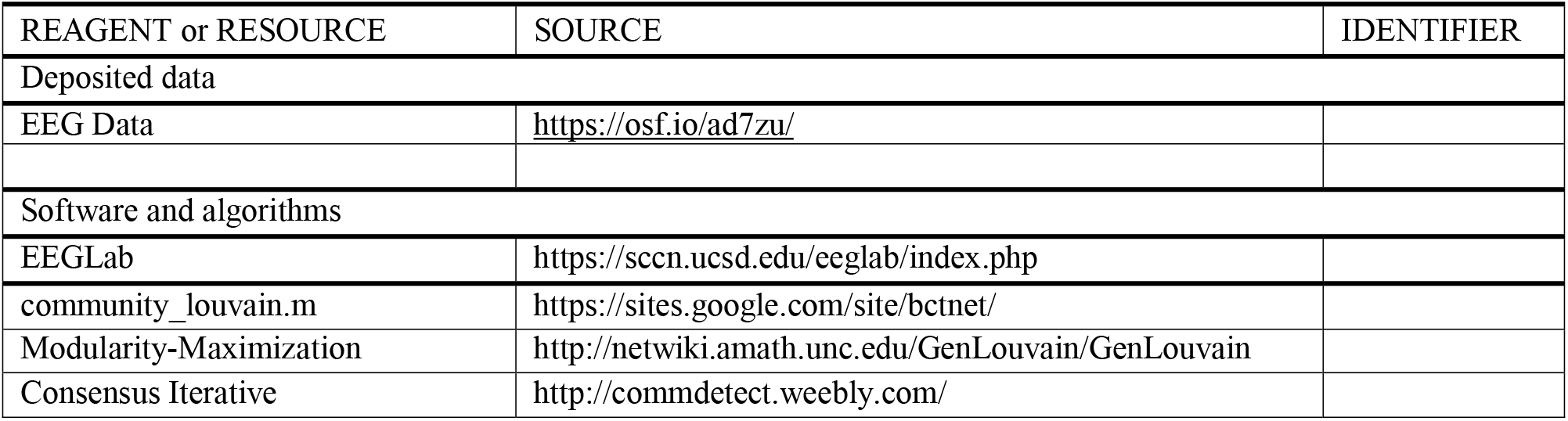
KEY RESOURCES TABLE.

### Participants

Twenty-eight subjects (twelve men; age range 18-30) took part in the experiment (three excluded due to not completing the full experiment). All subjects were recruited from the University of California, Santa Cruz (UCSC) for course credit. All of them had normal or corrected-to-normal vision, no history of psychiatric illness or head injury, reported no color-blindness They provided written informed consent. All procedures were approved by the ethical review board of UCSC.

### Stimulus and Apparatus

Stimuli were presented on a black background on a 53.4 cm electrically-shielded VPixxEEG monitor with a viewing distance of approximately 69.5 cm. The monitor operates at a refresh rate of 120 Hz with a resolution of 1920×1080. Stimulus presentation and behavioral data collection were controlled by Psychtoolbox-3 running in MATLAB. The stimuli consisted of 150 white dots presented within a 5-degree circular aperture centered on fixation. For each stimulus, a proportion (1%, 4.5%, 8%, 12%, 25%, or 40%) of the dots were randomly selected on each frame to be displaced by a fixed distance of .5 degrees in either the left or right direction on the following frame. The rest of the dots were placed randomly and independently within the circular aperture. A small red fixation dot was presented at the center of the stimulus throughout the entire trial.

### Procedure

Participants first completed a practice block with 180 trials, with an equal number of trials for each coherence level. Auditory feedback was presented only on the practice trials. Participants were able to proceed to the main task once they reached over 80% accuracy at the highest coherence level. Participants who failed to reach these criteria in the very first practice block performed more practice blocks until they met the criteria.

Each trial began with a red dot presented in the center of the screen for a random inter-trial interval between 1000 and 1500 ms. The dots appeared for 300 ms, with the red dot remaining on the screen. Participants gave their choice and confidence response (1-4; 1 = guessing, 4 = highly confident) with a single button press using their left hand (from pinky finger to index finger) respectively on keys ‘A’, ‘S’, ‘D’, and ‘F’ to indicate left motion with the confidence rating from 4-1. and their right hand (from index finger to pinky finger) respectively on key ‘J’, ‘K’, ‘L’, and ‘;” representing right motion with the confidence rating from 1-4. Participants had unlimited time to respond.

Each participant completed 1080 trials in total, consisting of 180 trials with each motion coherence level. The trials were presented in six blocks with 180 trials in each block. Motion direction and coherence level varied independently and randomly on a trial-by-trial basis.

### EEG recording and analysis

EEG was acquired from 64 active electrodes (BrainVision ActiChamp), with each impedance kept below 30kΩ. Data was digitized at 1000 Hz and FCz was used as the online reference. EEG was processed offline using custom scripts in MATLAB (version R2019b) and with EEGLAB toolbox^46^. Recordings were down-sampled to 500 Hz and high-pass filtered at 0.1 Hz using a zero-phase, Hamming-windowed FIR filter. Data were re-referenced offline to the average of all electrodes. EEG data were segmented into epochs centered on stimulus onset using a time window of -2000 to 2000 ms. Individual trials were rejected if any scalp channel exceeded 100 μV at any time during the interval extending from -500 to 500 ms relative to the stimulus onset. On average, 209 trials were rejected for each participant. These trials were not involved in the analysis of behavioral data. Noisy channels were spherically interpolated and independent components analysis was performed to remove components reflecting eye-blinks or eye movements. A pre-stimulus baseline of -200 to 0 ms was subtracted from each trial.

### Modularity-maximization based clustering analysis

EEG data for each trial were pooled from all 25 participants resulting in 20983 trials. The trials were pooled among participants to ensure consistency in clustering correspondence. A trial-by-trial similarity matrix was created using the Pearson Correlation between trials calculated from stimulus onset (0 ms) to 500 ms after across 63 sensors, corresponding to 15813 data points per trial. The correlation period used was limited to an interval from stimulus onset (0 ms) to 500 ms post-stimulus because this period encompassed the time window associated with sensory processing. We note that we also performed a control analysis where we extended the time window to encompass the period from 0 to 1000 ms and still found similar results (**Figure S4**).

The similarity between trials was calculated using Pearson correlation because this is a commonly used measure to estimate the similarity ^19^. The clustering utilized both spatial and temporal activity. The spatial data is represented from all EEG sensors (63 channels) and temporal activity is from stimulus onset to 500 ms post-stimulus sampled every 2 ms for a total of 63 (channels) x 251 (timepoints) = 15813 data points from each trial. This many data points raise the possibility that clustering might be noisy. To ensure maximum signal quality, we eliminated bad trials (see above) and validated the robustness of our clustering results using cross-validation. Critically, trials from both subtypes were present in all subjects suggesting that the brain activity identified is consistent and robust a consistent pattern of activity.

The clustering procedure was conducted in the same manner as previously described in Nakuci *et al* ^19^. Modularity-maximization was implemented with the Generalized Louvain algorithm^47^ to cluster the similarity matrix^21^. We opted for modularity-maximization because unlike other clustering methods, such as *k*-means, modularity-maximization does not require the number of clusters to be *a priori* defined. Nevertheless, it is possible for this type of analysis to be conducted with other clustering methods^48,49^. Conceptually, our analysis is similar to the microstate analyses which aim to identify the brain state at each time point^22^. Our framework and analysis extend the concept of a microstate to states that last for hundreds of milliseconds.

To avoid suboptimal partition results that can depend on the initial seeding, the clustering was repeated 100x, and the final partition was based on the consensus across iterations using *consensus_iterative.m*^50^. Specifically, consensus clustering identifies a single representative partition from the set of 100 iterations. This process involves the creation of a thresholded nodal association matrix which describes how often nodes are placed in the same cluster. The representative partition is then obtained by using a Generalized Louvain algorithm to identify clusters within the thresholded nodal association matrix^50^.

### Identification of event-related potentials (ERPs)

ERPs for each subtype were generated by averaging all trials within the corresponding clusters across all sensors. Similarly, for experimental – motion direction and coherence levels – and behavioral factors – accuracy, response times, and confidence – ERPs were derived by using trials for each factor, respectively. Motion direction and motion coherence ERPs were derived by averaging left/right moving trials and each of the 6 motion coherence levels. ERPs for the behavioral factor were derived by averaging correct/incorrect trials. For response times trials were first separated into fast/slow bins if they were higher or lower than the median response time, then trials in each bin were averaged to obtain ERPs representing fast and slow responses, respectively. In the same manner for confidence, trials were first separated into high/low confidence bin based on if the confidence for a trial was above or below the average confidence values and then all trials within the high or low confidence bin were averaged, respectively. The similarity between the ERPs from Subtype 1 and Subtype 2 to experimentally and behaviorally defined ERPs was based on estimating the Pearson correlation across all sensors from 0 to 1000 ms.

### Topographic analysis of variance

To identify the time periods in which the ERP topographies differed between subtypes, we used a topographic analysis of variance (TANOVA)^22^. TANOVA calculates the topographical similarity at each time period between the subtype-derived ERPs to ERPs derived from permuted labels. For each period, the labels were randomized 1000x. Time periods in which the empirical topographic similarity was less than the similarity derived from randomized labels with p < 0.05 following FDR correction were considered significant.

### Drift-diffusion modeling

We fit the diffusion model^34^ to the data using the hierarchical drift-diffusion model (HDDM) python package^51^. The model allowed the drift rate parameter to vary across motion coherence levels, but the other parameters, boundary and non-decision time, were fixed. The boundary and non-decision time parameters were estimated as part of the model fitting. Model fitting was conducted separately for each trial subtype.

### Reliability and robustness analysis

A 5-fold cross-validation analysis using Support Vector Machine (SVM) classifier was performed using MATLAB’s *fitcsvm.m.* Trials were randomly separated into 5 bins containing 20% of trials. The SVM classifier was trained on EEG data from 4 of the bins (80% of trials) and tested on the remaining bin (20% of the trials). This procedure was repeated until each bin was tested. Second, we replicated the clustering analysis with a longer time window (1000 ms).

### Statistical analysis and software

All data processing and statistical analyses were performed in MATLAB 2019b.

### Data and code availability

All data and code are available at https://osf.io/ad7zu/

## Acknowledgments

This work was supported by the National Institutes of Health (grant R01MH119189 to DR) and the Office of Naval Research (grant N00014-20-1-2622 to DR).

