## Supplemental Information for "Brain signatures indexing variation in internal processing during perceptual decision-making"

**This PDF file includes:**

Figs. S1 to S4

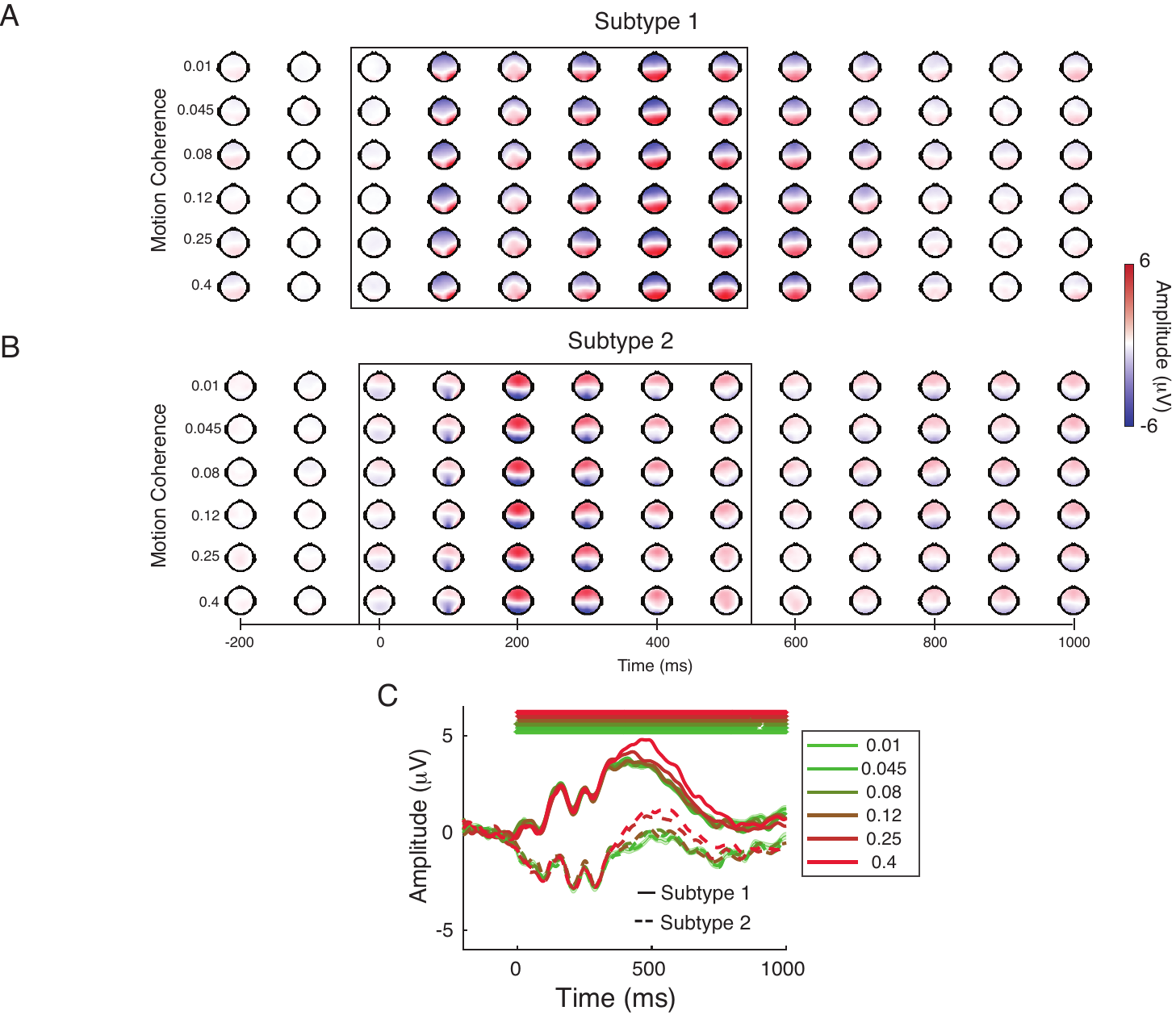

**Figure S1.** ERP topography of each motion coherence level for A) Subtype 1 and B) Subtype 2. The black box highlights the period used for clustering. C) ERP activity from the centro-parietal sensor per motion coherence level across subtypes. Each waveform shows the mean and standardized error of the mean ^30^. Statistical testing was conducted using independent samples t-tests, and FDR corrected for multiple comparisons. Statistically significant differences in amplitude are marked at the top of each panel.

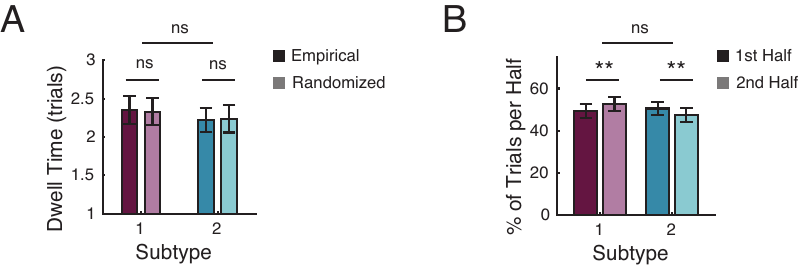

**Figure S2.** Dwell time and half-split comparison. A) The dwell time measuring the average number of consecutive trials between Subtype 1 or Subtype 2. B) The proportion of trials labeled either as Subtype 1 or Subtype 2 in the first and second half of the experiment. ** p < 0.01, ns: p > 0.05.

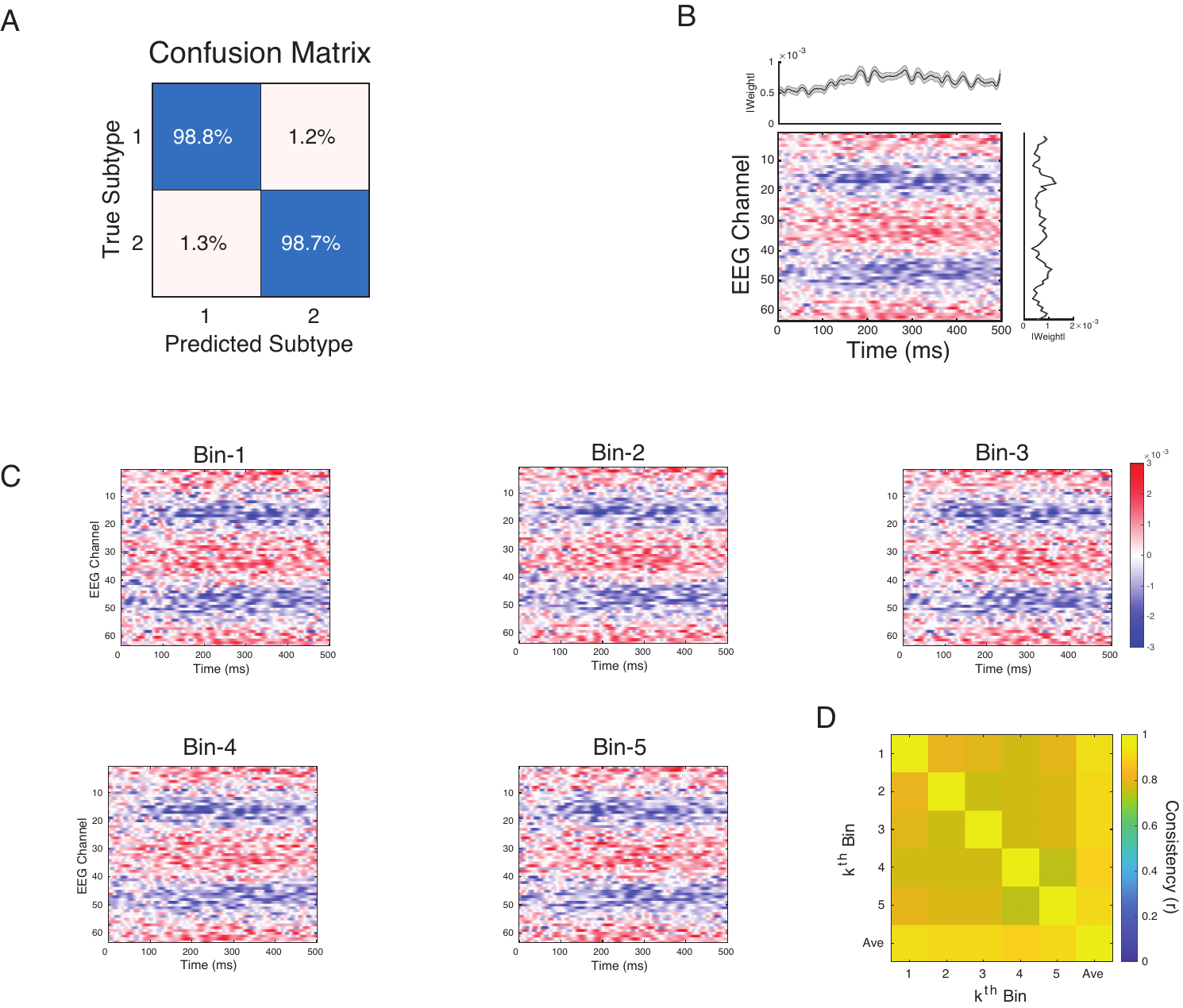

**Figure S3.** SVM classification weights. A) Cross-validation confusion matrix showing SVM classification accuracy. B) SVM weights present the average weights from the 5-fold cross-validation analysis. The average SVM weight across time (*top insert*) and EEG channels (*right insert*)*,* respectively. C) SVM weights indicating how important the EEG activity from a channel at a specific time point was for each of the 5-fold cross-validation bins. D) Consistency in the SVM weights across each of bin and to the overall average. Consistency was calculated using Pearson correlation.

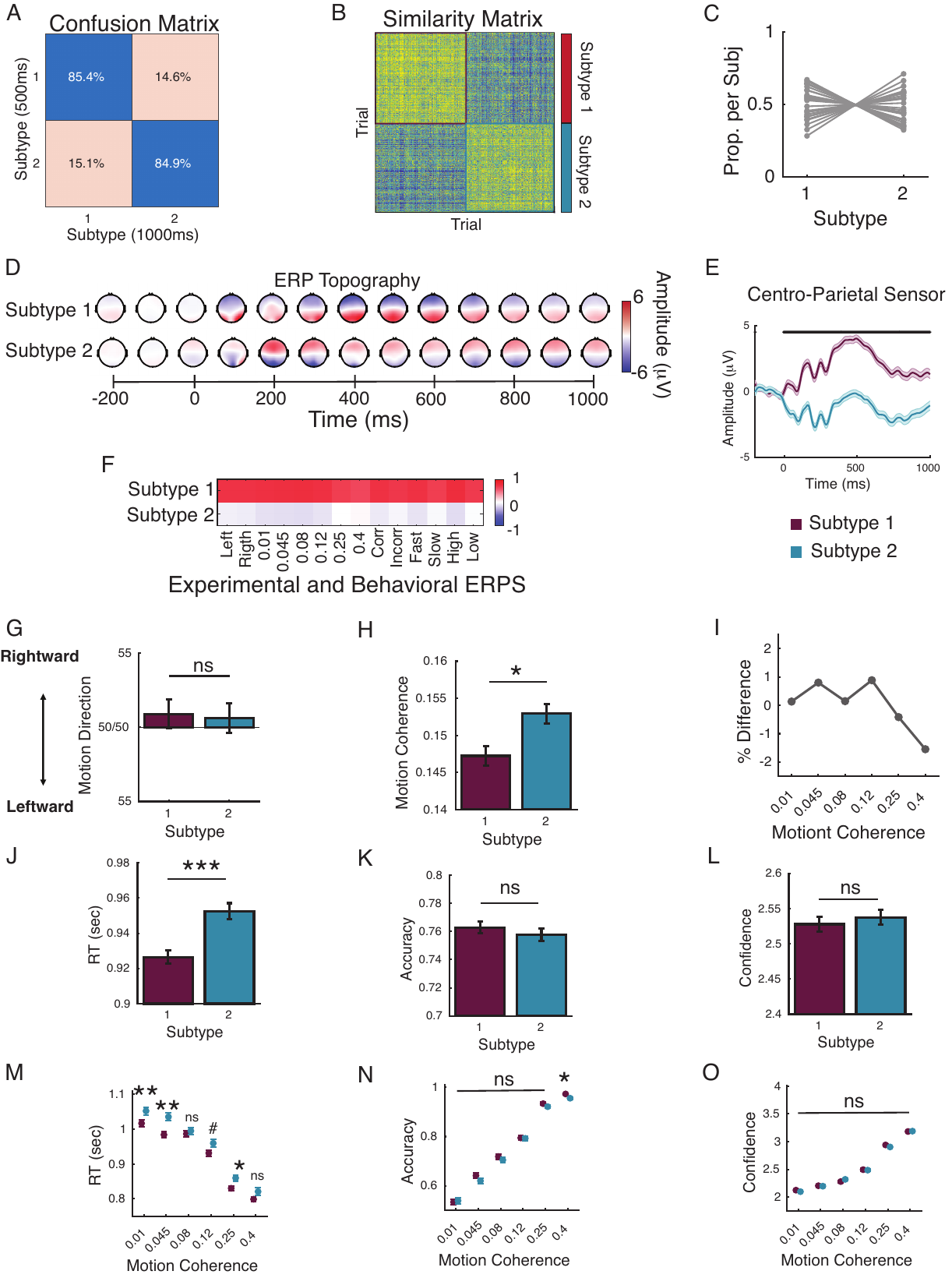

**Figure S4.** Subtypes of individual trials in motion perception task from 1000 ms window. A) Confusion matrix between trial subtype labels between the 500 ms and 1000 ms window used to calculate the similarity. B) Similar to the results in the manuscript, modularity-maximization based clustering identified two subtypes of trials, Subtype 1 and Subtype 2, when using the similarity was calculates from 0 ms to 1000 ms after the stimulus. C) The proportion of trials in each subject classified as either subtype 1 or 2. D) ERP topographies of Subtype 1 and Subtype 2 from stimulus onset (0 ms) to 1000 ms after the stimulus. E) ERP activity from the centro-parietal sensor per subtype (mean ± sme) ^30^. Statistical testing was conducted using an independent samples t-tests and FDR corrected for multiple comparisons. Statistically significant differences in amplitude are marked at the top of each panel. F) Topographical similarity between ERPs derived from each subtype with ERPs derived from experimental – motion direction (Left/Right), motion coherence (0.01, 0.045, 0.08, 0.12, 0.25 0.4) – and behavior factors – Correct/Incorrect response, Fast/Slow response time, High/Low confidence. Corresponding differences in G) Motion Direction and H) Motion Coherence level between subtypes. I) Differences in the percent of trials between subtypes per motion coherence level. Differences in J) Response Times, K) Accuracy, and L) Confidence between subtypes, respectively. Error bars show the mean ± sem. Behavioral performance differences between subtypes per motion coherence level in M) Response Times, N) Accuracy, and O) Confidence between subtypes, respectively (mean ± sem). Statistical testing was conducted using an independent samples t-tests and FDR corrected for multiple comparisons. *** p < 0.001, FDR; # p < 0.05, uncorrected
